# Dual activity of PNGM-1, a metallo-β-lactamase and tRNase Z, pinpoints the evolutionary origin of subclass B3 metallo-β-lactamases

**DOI:** 10.1101/575373

**Authors:** Jung Hun Lee, Masayuki Takahashi, Jeong Ho Jeon, Lin-Woo Kang, Mineaki Seki, Kwang Seung Park, Myoung-Ki Hong, Yoon Sik Park, Tae Yeong Kim, Asad Mustafa Karim, Jung-Hyun Lee, Masayuki Nashimoto, Sang Hee Lee

## Abstract

Antibiotic resistance is a steadily increasing global problem which could lead to a fundamental upheaval in clinical care with the potential to return us to the pre-antibiotic era^1-4^. The production of β-lactamases, a group of enzymes that confer antibiotic resistance in Gram-negative bacteria, is now one of the major barriers in treating Gram-negative infections^5^. β-Lactamases are classified according to their catalytic mechanisms into serine β-lactamases and metallo-β-lactamases^6,7^. There are functional and structural similarities between serine β-lactamases and penicillin-binding proteins, and so serine β-lactamases are thought to have evolved from a penicillin-binding protein^7,8^. Given the functional and structural differences between serine β-lactamases and metallo-β-lactamases, metallo-β-lactamases are thought to have evolved from a protein other than a penicillin-binding protein, but to date this ancestor remains unknown^8-11^. We discovered PNGM-1, the first subclass B3 metallo-β-lactamase, in deep-sea sediments that predate the antibiotic era^12^. Here we discover the dual activity of PNGM-1, pinpointing the evolutionary origin of subclass B3 metallo-β-lactamases. Phylogenetic analysis suggested that PNGM-1 could yield insights into the evolutionary origin of subclass B3 metallo-β-lactamases. We reveal the structural similarities between tRNase Zs and PNGM-1, which prompted us to investigate their evolutionary relationship and the possibility of them possessing dual enzymatic activities. We demonstrate that PNGM-1 has dual activity with both true metallo-β-lactamase and tRNase Z activity, suggesting that PNGM-1 is thought to have evolved from a tRNase Z. We also show kinetic and structural comparisons between PNGM-1 and other proteins including subclass B3 metallo-β-lactamases and tRNase Zs. These comparisons revealed that the B3 metallo-β-lactamase activity of PNGM-1 is a promiscuous activity and subclass B3 metallo-β-lactamases are thought to have evolved through PNGM-1 activity. Our work provides a foundation for the evolution of tRNase Z into subclass B3 metallo-β-lactamases through the dual activity of PNGM-1.

## Introduction

β-Lactam antibiotics are one of the most successful drugs used for the treatment of bacterial infections and represent roughly 65% of the total world market for antibiotics^13^. Therefore, resistance to β-lactam antibiotics is one of the most serious problems associated with Gram-negative bacterial infections^5^. β-Lactamases are produced by various bacteria conferring them resistant to β-lactam antibiotics such as penicillins, cephalosporins, monobactams, and/or carbapenems (the last resort drugs for treating bacterial infections)^9,14^. β-Lactamases are divided into four classes, A–D. The enzymes belonging to classes A, C, and D, are serine β-lactamases, which are thought to originate from a penicillin-binding protein^7,8^. Class B enzymes, which are further divided into three subclasses, B1–B3^6^, are metallo-β-lactamases (MBLs) that hydrolyse almost all β-lactam antibiotics, including carbapenems, thereby representing a critical antibiotic resistant threat^10,11^. The evolutionary origin of MBLs is currently unknown.

Subclass B1 and B3 MBLs require two zinc ions for maximum β-lactamase activity, whereas the subclass B2 MBLs require only one^15^. Recently, we discovered a novel B3 MBL named PNGM-1 (<u>P</u>apua <u>N</u>ew <u>G</u>uinea <u>M</u>etallo-β-lactamase), which was the first B3 enzyme obtained from a functional and bacterial metagenomic library of deep-sea sediments from the Edison seamount, which existed prior to the antibiotic era^12,16^.

MBLs belong to an ancient MBL superfamily^17,18^. Members of this superfamily are widespread across the three biological domains (Eukarya, Archaea and Bacteria) and engage in many diverse functions. Members of this superfamily include MBLs, a Zn-dependent hydrolase, lactonase, N-acyl homoserine lactone hydrolase, 4-pyridoxolactonase, alkylsulfatase, sulfur dioxygenase, a teichoic acid phosphorylcholine esterase, a methyl parathion hydrolase, a human glyoxalase II, β-CASP metallo-β-lactamase family nuclease, tRNase Zs from all three domains and so forth^17,19-23^. Amino acid sequence identity between the members is low (less than 5%), but structural features, i.e., αββα-fold or MBL fold, and the unique metal binding motif (HXHXDH) are shared^20,21^.

A member of the MBL superfamily, tRNase Z, is a tRNA processing enzyme which removes the 3′ trailer from pre-tRNA^24,25^. Most tRNase Zs cleave pre-tRNA immediately downstream of a discriminator nucleotide (nt), onto which the CCA residues are added to produce mature tRNA. tRNase Zs are categorized into two groups, tRNase Z^S^ containing 300–400 amino acids and tRNase Z^L^ containing 800–900 amino acids. Bacteria and archaea have tRNase Z^S^ only, while eukaryotes have either tRNase Z^L^ only or both tRNase Z^S^ and tRNase Z^L^. tRNase Z (prokaryotic tRNase Z^S^) can cleave unstructured single-strand RNAs that are unrelated to pre-tRNA *in vitro*^26^.

The tRNase Z catalytic center is formed by the major five well conserved residues (His-48, His-50, Asp-52, His-53 and His-222 in the case of *T. maritima* tRNase Z) together with two zinc ions. Residues Asp-52 and His-222 are thought to directly contribute as donors during the catalytic proton transfer^27^.

In this paper, we found that in addition to β-lactamase activity, PNGM-1 possesses endoribonuclease (tRNase Z) activity on both pre-tRNA substrates and on small unstructured single-strand RNA substrates. Our functional, phylogenetic and structural analyses of PNGM-1 and their comparison with the properties of other MBLs (proteins containing the αββα-fold with true-β-lactamase activity) and structurally representative MBL fold proteins (proteins having αββα-fold without true-β-lactamase activity) of the MBL superfamily, reveal the evolution of tRNase Z to subclass B3 MBLs through the dual activity of PNGM-1.

## Materials and Methods

### Strains and plasmids

*Escherichia coli* BL21 (DE3) and plasmid-containing *E. coli* strains were used for all cloning and expression studies. The pET-28a(+)/His_6_-PNGM-1 plasmid has been described previously^12^. The strains and plasmids used in this study are listed in Supplementary Table S1.

### Construction of PNGM-1 mutants by site-directed mutagenesis

Site-directed mutagenesis was carried out using a QuikChange II Site-Directed Mutagenesis Kit (Stratagene, Agilent Technologies, Santa Clara, CA, USA) according to the manufacturer’s instructions. H91A, H93A, D95A, H96A and H257A mutants were generated, with the template (pET-28a(+)/His_6_-PNGM-1) and primer sequences listed in Supplementary Table S2. After verifying the DNA sequences, the plasmids, [pET-28a(+)/His_6_-PNGM-1(H91A), pET-28a(+)/His_6_-PNGM-1(H93A), pET-28a(+)/His_6_-PNGM-1(D95A), pET-28a(+)/His_6_-PNGM-1(H96A) and pET-28a(+)/His_6_-PNGM-1(H257A)], were individually transformed into *E. coli* BL21 (DE3) cells.

### Cloning of *bla*_AIM-1_, *bla*_GOB-18_, *bla*_FEZ-1_ and three tRNase Z genes

Six DNA templates encoding *bla*_AIM-1_ gene (GenBank ID: AM998375), *bla*_GOB-18_ gene (GenBank ID: DQ004496), *bla*_FEZ-1_ gene (GenBank ID: Y17896) and three tRNase Z genes from *Bacillus subtilis* (Bs-tRNase Z, GenBank ID: WP_101502431), *E. coli* (Ec-tRNase Z, GenBank ID: Q47012) and *T. maritima* (Tm-tRNase Z, GenBank ID: NP_228673), which are codon optimized for *E. coli*, were synthesized and purchased from IDT (Integrated DNA Technologies, Coralville, IA, USA). The DNA templates were amplified by PCR using suitable primer pairs (Supplementary Table S2). The amplified DNA and pET-30a(+) vector (Novagen, Madison, Wisconsin, USA) were double-digested with *Nde*I and *Xho*I, with digested DNA then ligated into the digested vector. After verifying the DNA sequences, the plasmids, pET-30a(+)/His_6_-*bla*_AIM-1_, pET-30a(+)/His_6_-*bla*_GOB-18_, pET-30a(+)/His_6_-*bla*_FEZ-1_, pET-30a(+)/His_6_-Bs-tRNase Z, pET-30a(+)/His_6_-Ec-tRNase Z plasmid and pET-30a(+)/His_6_-Tm-tRNase, were individually transformed into *E. coli* BL21 (DE3) cells.

### Preparation of PNGM-1, PNGM-1 mutants, AIM-1, GOB-18, FEZ-1 and tRNase Zs

Each of the histidine-tagged proteins, PNGM-1, five PNGM-1 mutants (H91A, H93A, D95A, H96A, and H257A), AIM-1, GOB-18, FEZ-1, Bs-tRNase Z, Ec-tRNase Z, Tm-tRNase Z and human Δ30 tRNase Z^L^ (lacking the N-terminal 30 amino acids) were prepared as previously described^12,24,28^.

### Steady-state kinetic analysis

Kinetic assays were conducted at 30°C with a Shimadzu UV-1650PC spectrophotometer (Shimadzu Corp.). β-Lactam hydrolysis was detected by monitoring the change in absorbance using the characteristic molar extinction coefficient of each substrate: cefoxitin (Δε_270nm_ = −8,380 M^-1^ cm^-1^); ceftazidime (Δε_265nm_ = −10,300 M^-1^ cm^-1^); cefotaxime (Δε_264nm_ = −7,250 M^-1^ cm^-1^); imipenem (Δε_278nm_ = −5,660 M^-1^ cm^-1^); meropenem (Δε_298nm_ = −9,530 M^-1^ cm^-1^); and ertapenem (Δε_295nm_ = −10,940 M^-1^ cm^-1^). The assays were conducted in 50 mM MES [2-(N-morpholino)ethanesulfonic acid] (pH 7.0) buffer which contained the enzyme, 100 μM ZnCl_2_ and 100 μg ml^−1^ bovine serum albumin. Steady-state kinetic constants were determined by fitting the initial rates (in triplicate) directly to the Michaelis-Menten equation using nonlinear regression with the program DynaFit^29^.

### Construction of the phylogenetic tree

Multiple sequence alignments of PNGM-1 with 82 representative enzymes of subclasses B1, B2 and B3 MBLs from the Beta-Lactamase DataBase (http://bldb.eu/) and structurally representative MBL fold proteins [a Zn-dependent hydrolase of *Thermotoga maritima* (Tm), another Zn-dependent hydrolase of *T. maritima* (Tm-1), lactonase from *T. maritima* (Tm-Lac), N-acyl homoserine lactone hydrolase from *Bacillus thuringiensis* (AiiA), 4-pyridoxolactonase from *Mesorhizobium japonicum* MAFF 303099 (PDLA), alkylsulfatase from *Pseudomonas aeruginosa* PAO1 (SdsA1), sulfur dioxygenase of *Arabidopsis thaliana* (ATSD), a teichoic acid phosphorylcholine esterase from *Streptococcus pneumoniae* (CbpE), a methyl parathion hydrolase from *Pseudomonas* sp. strain WBC-3 (Pah), a human glyoxalase II (Gox), β-CASP metallo-β-lactamase family nuclease from *Methanothermobacter thermautotrophicus* (MTH1203) and tRNase Zs]^20^ were performed with MAFFT (Multiple Alignment using Fast Fourier Transform) in the Geneious software (v11.1.5, Biomatters, http://www.geneious.com). The phylogenetic tree was constructed by the neighbor-joining method using the Molecular Evolutionary Genetics Analysis 6.0 software (MEGA, version 6.0)^30^. Accession numbers of all enzymes used for phylogenetic analysis are listed in Supplementary Table S3.

### Structure determination of PNGM-1

Crystallization and X-ray diffraction data collection of PNGM-1 was carried out as previously published^16^; native PNGM-1 crystals were obtained under the reservoir condition of 0.1 M Tris-HCl pH 7.0, 0.2 M MgCl_2_, 9% (w/v) PEG 8000 and selenomethionine--substituted (SeMet) PNGM-1 crystals were obtained under the reservoir condition of 0.1 M HEPES pH 7.5, 0.1 M CaCl_2_, 8.25% (w/v) PEG 3350, 1.32% (v/v) isopropanol. The native and SeMet PNGM-1 crystals were cryoprotected with the reservoir solution containing 20% (v/v) MPD (2-methyl-2,4-pentanediol). X-ray diffraction data were collected to 2.1 and 2.3 Å resolutions for the native and SeMet PNGM-1 crystals, respectively. All data were integrated and scaled using the *DENZO* and *SCALEPACK* crystallographic data-reduction routines^31^.

The PNGM-1 structure was solved by the SAD method using SeMet PNGM-1. The interpretable electron density was obtained at 2.3 Å resolution for the SeMet PNGM-1 data set using the single wavelength SAD protocol of AUTO-RICKSHAW, an automated crystal structure determination platform^32^ in the *P*21 space group. The native PNGM-1 structure was determined by molecular replacement with *MOLREP*^33^ using the SeMet PNGM-1 structure (Supplementary Table S4).

### Preparation of RNA and DNA substrates

Total RNA of the human multiple myeloma cell line KMM-1 was extracted with RNAiso Plus (Takara Bio, Shiga, Japan). A 24-nt RNA, usRNA1 (5′-GAGUGACUACCUCCAAGGCCCUUU-3′), a 22-nt RNA, usRNA9 (5′-GCCUGGCUGGCUCGGUGUAUUU-3′), and a 24-nt DNA, usDNA1 (5′-GAGTGACTACCTCCAAGGCCCTTT-3′), were chemically synthesized with a 5′-6-carboxyfluorescein dye and subsequently purified by high-performance liquid chromatography. These were obtained from JBioS (Saitama, Japan). An 84-nt human pre-tRNA^Arg^, (5′-GGGCCAGUGGCGCAAUGGAUAACGCGUCUGACUACGGAUCAGAAGAUUCCAGGUUCGACUCCUGGCUGGCUCGGUGUAAGCUUU-3′), was synthesized using T7 RNA polymerase (Takara Bio, Shiga, Japan) from a corresponding synthetic DNA template, and subsequently 5′-labeled with fluorescein as previously described^25^.

### *In vitro* RNA cleavage assay

Various RNA substrates and a negative control DNA substrate were mixed with PNGM-1 in 50 μl of a 0.2–1 mM MES (pH 6.8) buffer or with tRNase Z in 50 μl of 4 mM Tris-HCl (pH 8.0) buffer in the presence of 5–50 mM MgCl2 or MnCl_2_ at 25– 80 °C for 30–90 min. The reaction products for the total RNA of the human cell line KMM-1 were analysed by microfluidics-based automated electrophoresis with the Agilent 2100 Bioanalyzer (Agilent Technologies, CA, USA) according to the manufacturer’s protocol. The reaction products for the other substrates were separated on a 20% polyacrylamide 8 M urea gel or in the case of the reaction products for the DNA substrate with MnCl_2_, a 20% polyacrylamide native gel. After resolution of the reaction products, the gel was analysed with a Gel-Doc imager (Bio-Rad Laboratories, CA, USA).

### Data availability

Coordinates for the PNGM-1 atomic model have been deposited in the Protein Data Bank under the accession codes 6J4N (native) and 6JKW (SeMet).

## Results and Discussion

### β-Lactamase activity of PNGM-1 and PNGM-1 mutants

The subclass B3 MBL, PNGM-1, has recently been discovered from the metagenomic library of deep-sea sediments from the Edison seamount that predates the antibiotic era^12^. While PNGM-1 has low amino acid sequence identity with other B3 MBLs, it does possess the unique metal binding motif (H_116_XH_118_XD_120_H_121_; numbering according to the BBL scheme^7^) that is only present in subclass B3 MBLs. To test the true β-lactamase (MBL) activity of PNGM-1, four active-site mutants, two at the first zinc-binding site [H91A (H116A, numbering according to the BBL scheme^7^) and H93A (H118A)] and two at the second zinc-binding site [D95A (D120A) and H96A (H121A)], were generated by site-directed mutagenesis.

To investigate the β-lactam-hydrolysing activity of the four PNGM-1 mutants against β-lactam antibiotics, the catalytic properties of purified PNGM-1 and the four PNGM-1 mutants were assessed (Table 1). Kinetic assays revealed that all four mutants were unable to hydrolyze β-lactam antibiotics including extended-spectrum cephalosporins (ceftazidime and cefotaxime) and carbapenems (meropenem, imipenem and ertapenem). Substitution of residues in the metal binding motif had the most significant influence on the catalytic properties of PNGM-1 against cephalosporins and carbapenems. Therefore, this result indicated that PNGM-1 possesses true β-lactamase (MBL with its ability to hydrolyse carbapenems) activity.

**Table 1.**
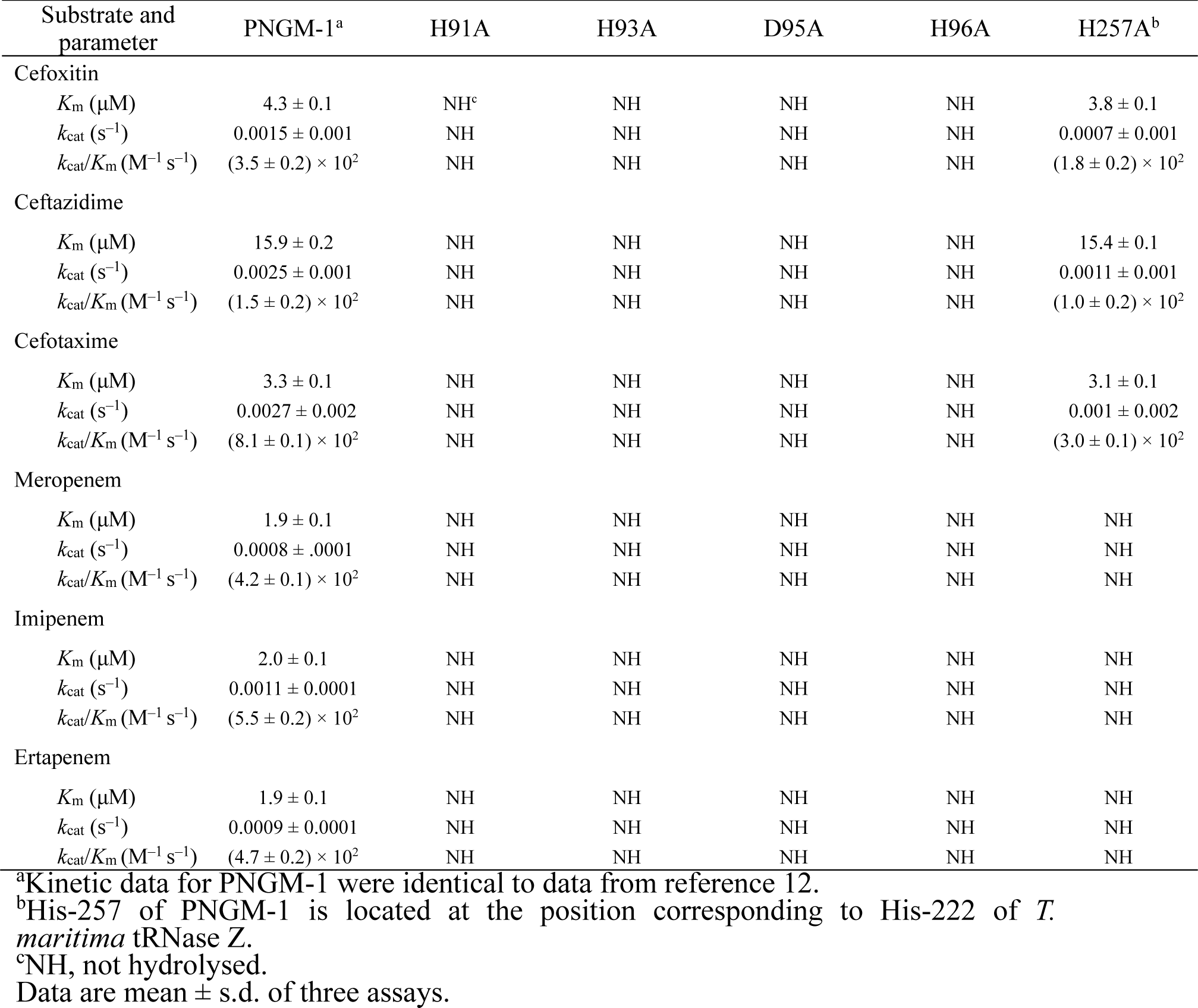
Kinetic parameters of PNGM-1 and five PNGM-1 mutants for various β-lactams

| Substrate and parameter | PNGM-1<sup>a</sup> | H91A | H93A | D95A | H96A | H257A<sup>b</sup> |
| --- | --- | --- | --- | --- | --- | --- |
| Cefoxitin |  |  |  |  |  |  |
| $K_m$ ($\mu\text{M}$) | $4.3 \pm 0.1$ | NH<sup>c</sup> | NH | NH | NH | $3.8 \pm 0.1$ |
| $k_{\text{cat}}$ ($\text{s}^{-1}$) | $0.0015 \pm 0.001$ | NH | NH | NH | NH | $0.0007 \pm 0.001$ |
| $k_{\text{cat}}/K_m$ ($\text{M}^{-1} \text{s}^{-1}$) | $(3.5 \pm 0.2) \times 10^2$ | NH | NH | NH | NH | $(1.8 \pm 0.2) \times 10^2$ |
| Ceftazidime |  |  |  |  |  |  |
| $K_m$ ($\mu\text{M}$) | $15.9 \pm 0.2$ | NH | NH | NH | NH | $15.4 \pm 0.1$ |
| $k_{\text{cat}}$ ($\text{s}^{-1}$) | $0.0025 \pm 0.001$ | NH | NH | NH | NH | $0.0011 \pm 0.001$ |
| $k_{\text{cat}}/K_m$ ($\text{M}^{-1} \text{s}^{-1}$) | $(1.5 \pm 0.2) \times 10^2$ | NH | NH | NH | NH | $(1.0 \pm 0.2) \times 10^2$ |
| Cefotaxime |  |  |  |  |  |  |
| $K_m$ ($\mu\text{M}$) | $3.3 \pm 0.1$ | NH | NH | NH | NH | $3.1 \pm 0.1$ |
| $k_{\text{cat}}$ ($\text{s}^{-1}$) | $0.0027 \pm 0.002$ | NH | NH | NH | NH | $0.001 \pm 0.002$ |
| $k_{\text{cat}}/K_m$ ($\text{M}^{-1} \text{s}^{-1}$) | $(8.1 \pm 0.1) \times 10^2$ | NH | NH | NH | NH | $(3.0 \pm 0.1) \times 10^2$ |
| Meropenem |  |  |  |  |  |  |
| $K_m$ ($\mu\text{M}$) | $1.9 \pm 0.1$ | NH | NH | NH | NH | NH |
| $k_{\text{cat}}$ ($\text{s}^{-1}$) | $0.0008 \pm .0001$ | NH | NH | NH | NH | NH |
| $k_{\text{cat}}/K_m$ ($\text{M}^{-1} \text{s}^{-1}$) | $(4.2 \pm 0.1) \times 10^2$ | NH | NH | NH | NH | NH |
| Imipenem |  |  |  |  |  |  |
| $K_m$ ($\mu\text{M}$) | $2.0 \pm 0.1$ | NH | NH | NH | NH | NH |
| $k_{\text{cat}}$ ($\text{s}^{-1}$) | $0.0011 \pm 0.0001$ | NH | NH | NH | NH | NH |
| $k_{\text{cat}}/K_m$ ($\text{M}^{-1} \text{s}^{-1}$) | $(5.5 \pm 0.2) \times 10^2$ | NH | NH | NH | NH | NH |
| Ertapenem |  |  |  |  |  |  |
| $K_m$ ($\mu\text{M}$) | $1.9 \pm 0.1$ | NH | NH | NH | NH | NH |
| $k_{\text{cat}}$ ($\text{s}^{-1}$) | $0.0009 \pm 0.0001$ | NH | NH | NH | NH | NH |
| $k_{\text{cat}}/K_m$ ($\text{M}^{-1} \text{s}^{-1}$) | $(4.7 \pm 0.2) \times 10^2$ | NH | NH | NH | NH | NH |

<sup>a</sup>Kinetic data for PNGM-1 were identical to data from reference 12.
<sup>b</sup>His-257 of PNGM-1 is located at the position corresponding to His-222 of *T. maritima* tRNase Z.
<sup>c</sup>NH, not hydrolysed.
Data are mean ± s.d. of three assays.

### The insight into the evolutionary origin of subclass B3 MBLs

The evolutionary origin of subclass B3 MBLs has not yet been identified. However, MBLs belong to an ancient MBL superfamily^17,18^ that evolved billions years ago^34,35^ and to date, there are more than 80 different types of MBLs in the Beat-Lactamase DataBase. A phylogenetic analysis was carried out with all MBL types, PNGM-1 (a unique B3 MBL) and 14 structurally representative MBL superfamily members (MBL fold proteins having no true β-lactamase activity: Tm, Tm-1, Tm-Lac, AiiA, PDLA, SdsA1, ATSD, CbpE, Pah, Gox, MTH1203, Bs-tRNase Z, Ec-tRNase Z and Tm-tRNase Z [Supplementary Table S3]). Our phylogenetic analysis revealed that (i) PNGM-1 was not grouped with subclass B3 MBLs but grouped with MBL fold proteins (Fig. 1a); (ii) PNGM-1 was closest to the tRNase Zs of the MBL fold proteins (Fig. 1a); and (iii) the HXHXDH motif was completely conserved in PNGM-1, the subclass B3 MBLs (AIM-1, GOB-18 and FEZ-1) and MBL fold proteins, but not in subclasses B1/B2 MBLs (Fig. 1b). These results suggest that PNGM-1 gives us an insight into the evolutionary origin of subclass B3 MBLs.

**Fig. 1.**
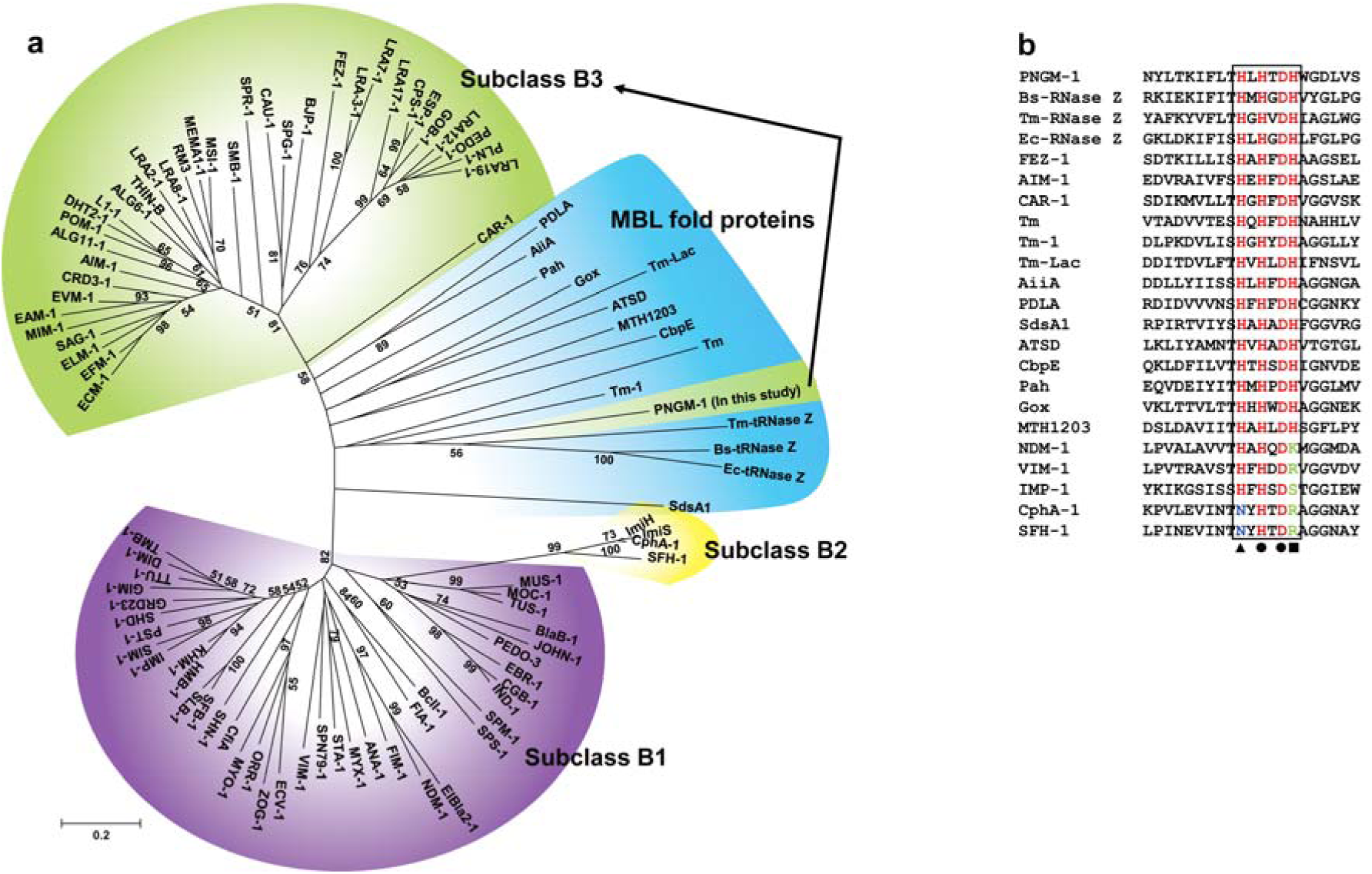
Characteristics of subclass B3 MBL PNGM-1. **a**, Neighbor-joining phylogenetic tree of PNGM-1 with all MBL types (B1, B2 and B3 MBL subclasses) and structurally representative enzymes of MBL fold proteins that have diverse functions but no β-lactamase activity. Only bootstrap values higher than 50% are shown. Bar, 0.2 substitutions per amino acid site. **b**, Multiple sequence alignment of PNGM-1 with representative sequences of B1, B2 and B3 MBLs and MBL fold proteins. The unique metal-binding motif, HXHXDH, is well conserved in subclass B3 MBLs including PNGM-1 and MBL fold proteins, and is indicated with a box. The closed triangle shows the difference between subclass B2 (CphA and SFH-1) and others, with the differing residues highlighted in blue, while the closed square shows the difference between B1 and B2 MBLs (NDM-1, VIM-1, IMP-1, CphA and SFH-1) and others, with the differing residues highlighted in green. The closed circles highlight residues which are conserved in all MBLs and MBL fold proteins. Accession numbers of all enzymes used for phylogenetic analysis are listed in Supplementary Table S3. Experiments in **a** and **b** were repeated at least three times and were reproducible.

### Structural similarity between PNGM-1 and tRNase Z

To examine the evolutionary relationship between PNGM-1 and tRNase Z, we compared the PNGM-1 structure with those of structurally representative tRNase Zs from *B. subtilis, E. coli* and *T. maritima* (Fig. 2). All the structures have the characteristic αββα-fold of the MBL superfamily with the unique metal binding motif. All structures had the characteristic αββα-fold of the MBL superfamily with the unique metal binding motif. The core of the PNGM-1 monomer is well superimposed on the tRNase Z monomers of Bs-tRNase Z, Ec-tRNase Z, and Tm-tRNase Z, with root-mean-square-deviations of 6.24 Å (168 α-carbons), 3.82 Å (170 α-carbons), and 5.76 Å (187 α-carbons), respectively.

**Fig. 2.**
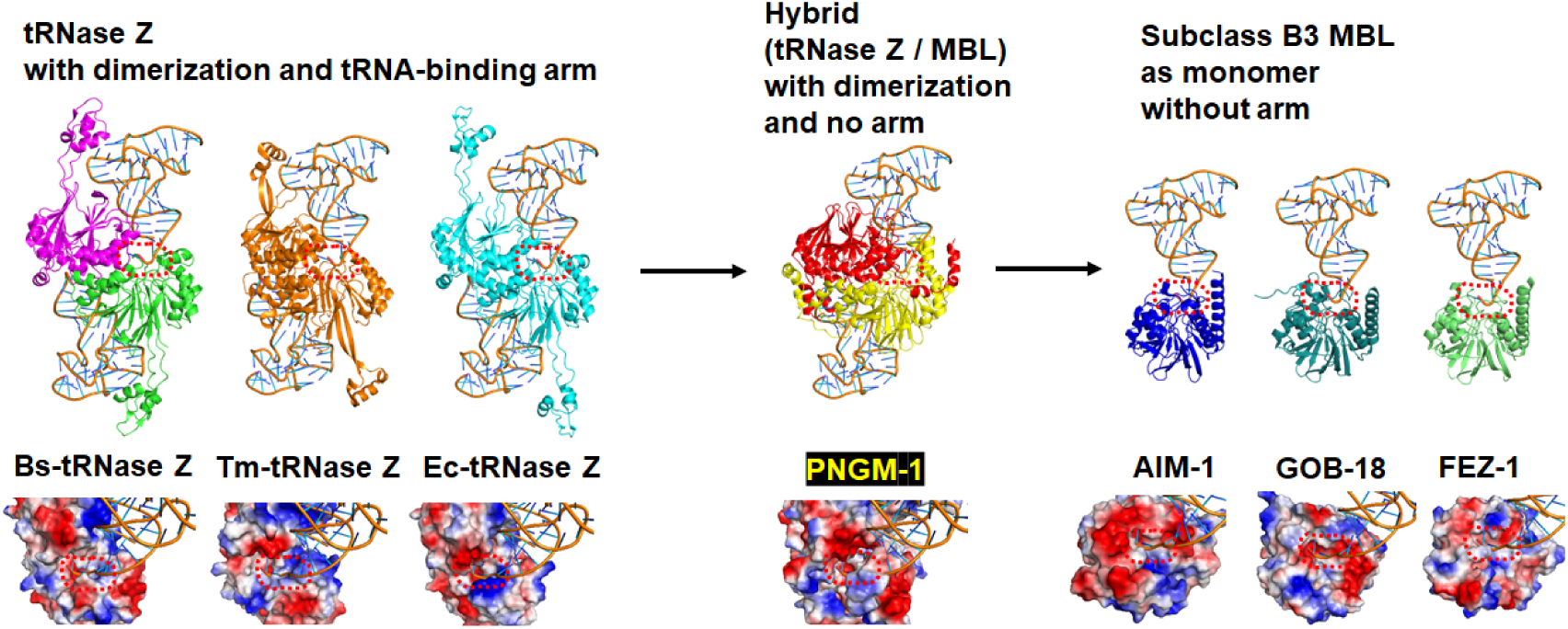
Gradual structural changes of tRNase Zs, PNGM-1 and subclass B3 MBLs. The structure of PNGM-1 [subunit A (red) and B (yellow), PDB entry 6J4N] was compared with those of tRNase Z from *B. subtilis* [Bs-tRNase Z (purple/green), PDB entry 4GCW], *T. maritima* [Tm-tRNase Z (orange/orange), PDB entry 2E7Y] and *E. coli* [Ec-tRNase Z (cyan/cyan), PDB entry 2CBN] and subclass B3 MBLs from *P. aeruginosa* [AIM-1 (blue), PDB entry 4AWZ], *Elizabethkingia meningoseptica* [GOB-18 (pale green), PDB entry 5K0W] and *Legionella gormanii* [FEZ-1 (lime), PDB entry 5W90]. The tRNA-bound Bs-tRNase Z structure was superimposed on the structures of the enzyme compared and the superimposed tRNA was presented with the enzymes (top row). In all structures, the 3’ end position of superimposed tRNA was bound in the substrate-binding pocket (bottom row). The substrate-binding pocket is marked with a red dashed line.

Structures of all tRNase Z enzymes showing dimerization is essential for tRNA binding and tRNase Z activity; tRNA is mostly bound to subunit A with the 3’ end bound to the active site of subunit B (Fig. 2). PNGM-1 structure also maintains the dimerization interactions and the dimer interface of PNGM-1 is strictly conserved to that of tRNase Z. When the tRNA-bound Bs-tRNase Z dimer was superimposed on the PNGM-1 dimer, tRNA can be bound to PNGM-1 subunit A without major hindrance and the 3’ end (to be processed) of tRNA is well positioned in the active site of subunit B (Figs. 2 and 3). The structurally representative subclass B3 MBLs, AIM-1, GOB-18 and FEZ-1, exist as monomers due to the presence of bulge-like structures at the proposed dimer interface which prevents dimerization due to steric hindrance (Figs. 2 and 3). tRNase Z has a conserved tRNA-binding arm, which is lost in PNGM-1 and subclass B3 MBLs (Fig. 2). These structural similarities between tRNase Zs and PNGM-1 prompts us to investigate the possibility of PNGM-1 possessing tRNase Z activity.

**Fig. 3.**
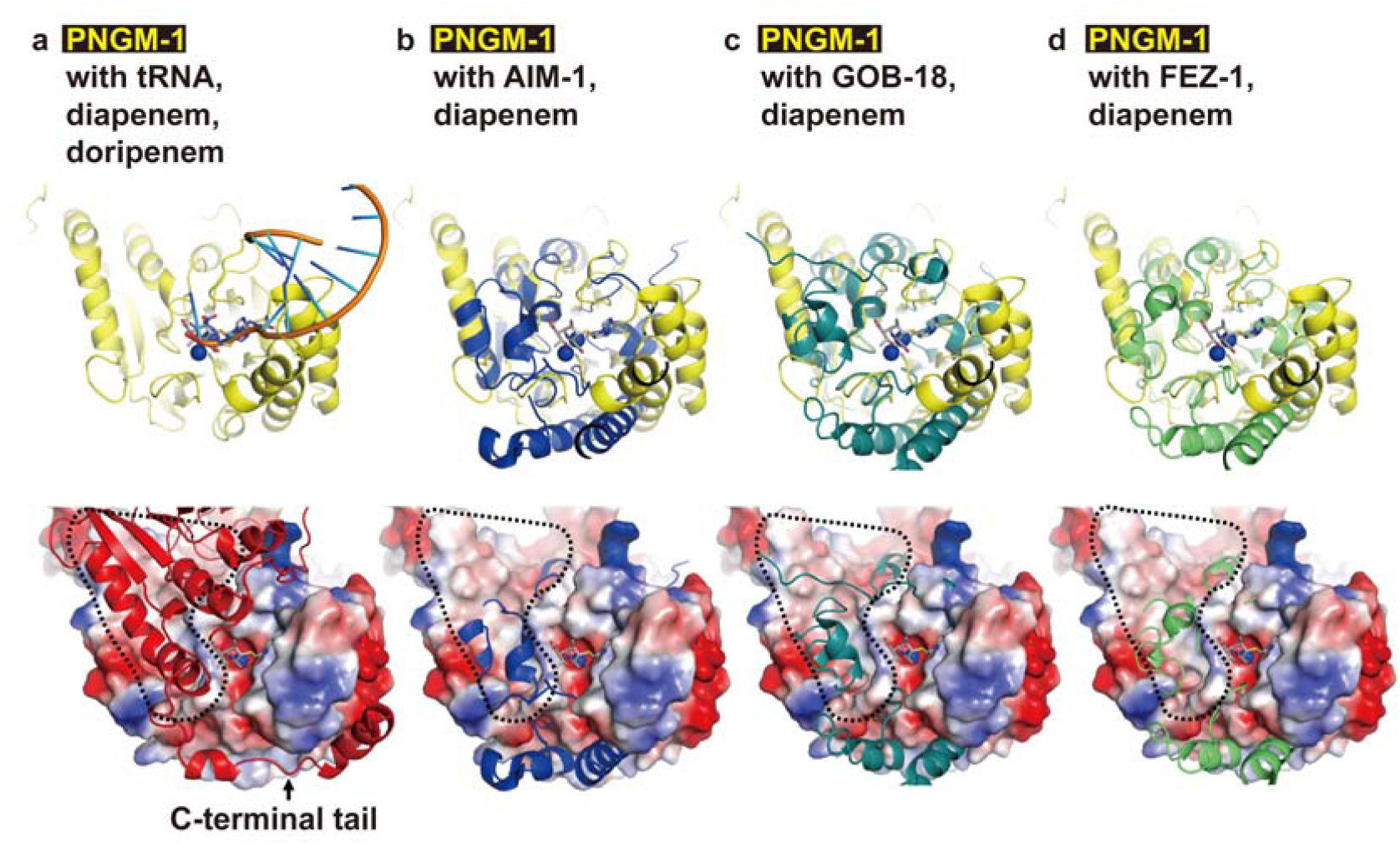
Structural comparison of PNGM-1 with subclass B3 MBLs. The structure of PNGM-1 (yellow) was superimposed with those of tRNA-bound Bs-tRNase Z (orange, PDB entry 4GCW), diapenem-bound CphA (grey, PDB entry 1X8I), doripenem-bound SMB-1 (pale purple, PDB entry 5B15), AIM-1 (blue, PDB entry 4AWZ), GOB-18 (pale green, PDB entry 5K0W)) and FEZ-1 (lime, PDB entry 5W90) (top row). **a**, PNGM-1 structure with superimposed tRNA, diapenem and doripenem. **b**, PNGM-1 structure with superimposed AIM-1 and diapenem. **c**, PNGM-1 structure with superimposed GOB-18 and diapenem. **d**, PNGM-1 structure with superimposed FEZ-1 and diapenem. The surface electrostatic potential of PNGM-1 (subunit A) is shown (bottom row). Structures of PNGM-1 (subunit B, red), AIM-1, GOB-18 and FEZ-1 are shown in cartoon representation. The main dimerization surface of PNGM-1 is marked with a black dashed line.

### PNGM-1 possesses RNase activity

To test whether PNGM-1 possesses RNase activity, we examined whether PNGM-1 can degrade total RNA isolated from the human cell line KMM-1. Total RNA was incubated with 15 μM PNGM-1 in the presence of 10 mM Mg^2+^ or Mn^2+^ at 37 °C for 60 min, and analysed by microfluidics-based automated electrophoresis. Both 18S and 28S rRNAs were degraded to smaller RNAs, and RNA species (∼100 nt) accumulated under both conditions (Fig. 4). The RNA integrity numbers in the assays with Mg^2+^ and Mn^2+^ decreased by 25% and 18%, respectively. These results suggest that PNGM-1 possesses a ribonuclease activity.

**Fig. 4.**
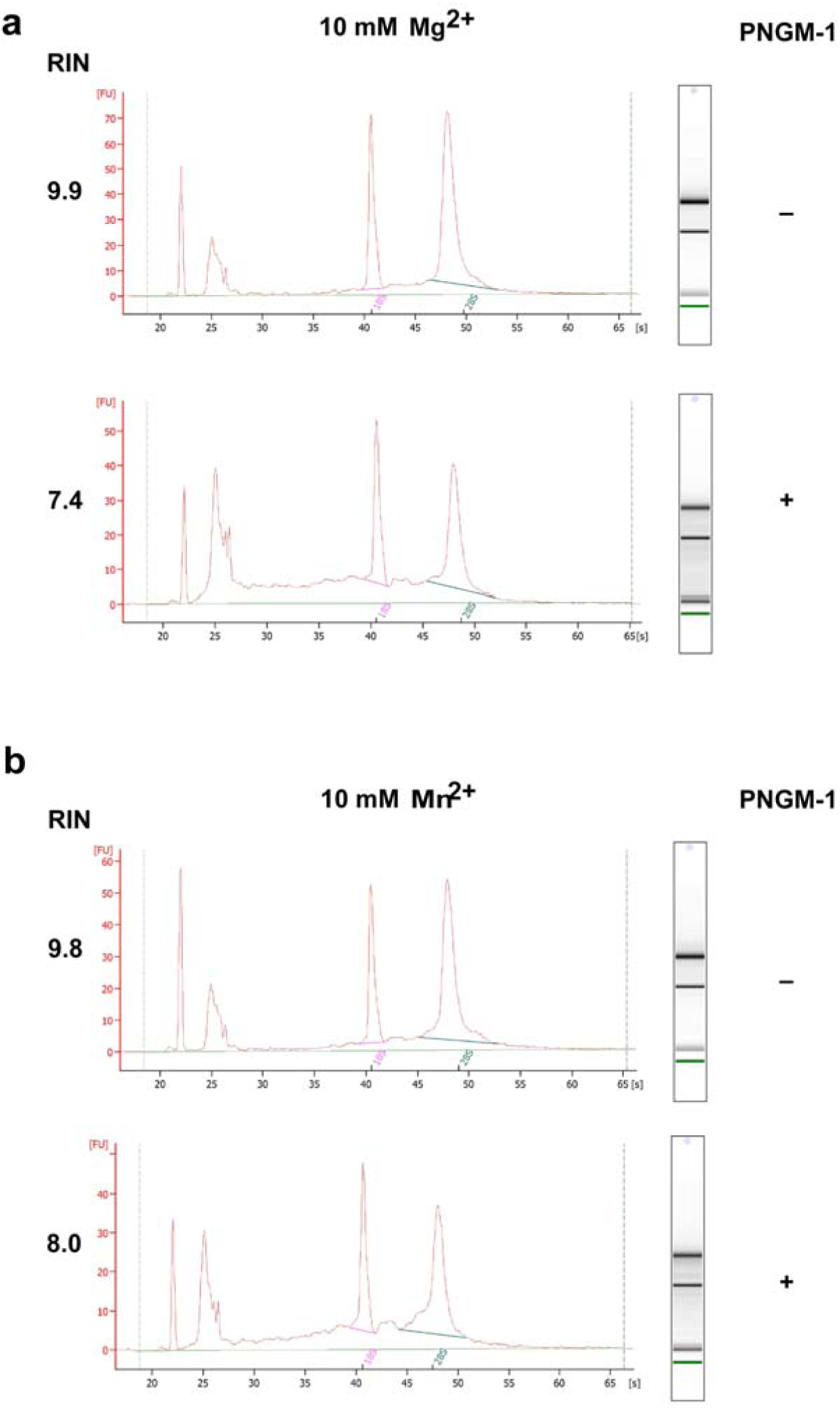
*In vitro* assays for RNase activity of PNGM-1 on total human cell RNA. **a**, **b**, Total RNA (0.3 μg) of the human cell line KMM-1 was incubated with or without 15 μM PNGM-1 in the presence of 10 mM MgCl_2_ (**a**) or MnCl_2_ (**b)**, at 37 °C for 60 min, and analysed by microfluidics-based automated electrophoresis. An electropherogram and a gel image for each sample are shown. The peaks around 22 and 25 seconds denote the 25-nt marker (green line on the gel image) and approximately 100-nt RNAs including tRNA, 5S rRNA and 5.8S rRNA, respectively. 18S, 18S rRNA (pink line); 28S, 28S rRNA (blue line); FU, arbitrary fluorescence units; RIN, RNA integrity number. Experiments in **a** and **b** were repeated at least three times and were reproducible.

### PNGM-1 can cleave unstructured RNAs endoribonucleolytically

Next, we examined PNGM-1 for RNase activity on two small RNA substrates. The unstructured RNAs, usRNA1 (24 nt) and usRNA9 (22 nt), were used as they are cleaved by tRNase Z. An unstructured 24-nt DNA (usDNA1), corresponding to usRNA1 was used as a negative control substrate. These three substrates, which were 5′-labeled with 6-carboxyfluorescein, were incubated with 15 μM PNGM-1 in the presence of 10 mM Mg^2+^ at 50 °C for 30–90 min, and the products were analysed by denaturing polyacrylamide gel electrophoresis. Fragments of various sizes were generated from usRNA1 and usRNA9 but not from usDNA1 (Fig. 5a). The pattern of generated RNA fragments suggest that PNGM-1 possesses an endoribonuclease activity, not an exoribonuclease activity, and has no DNase activity.

**Fig. 5.**
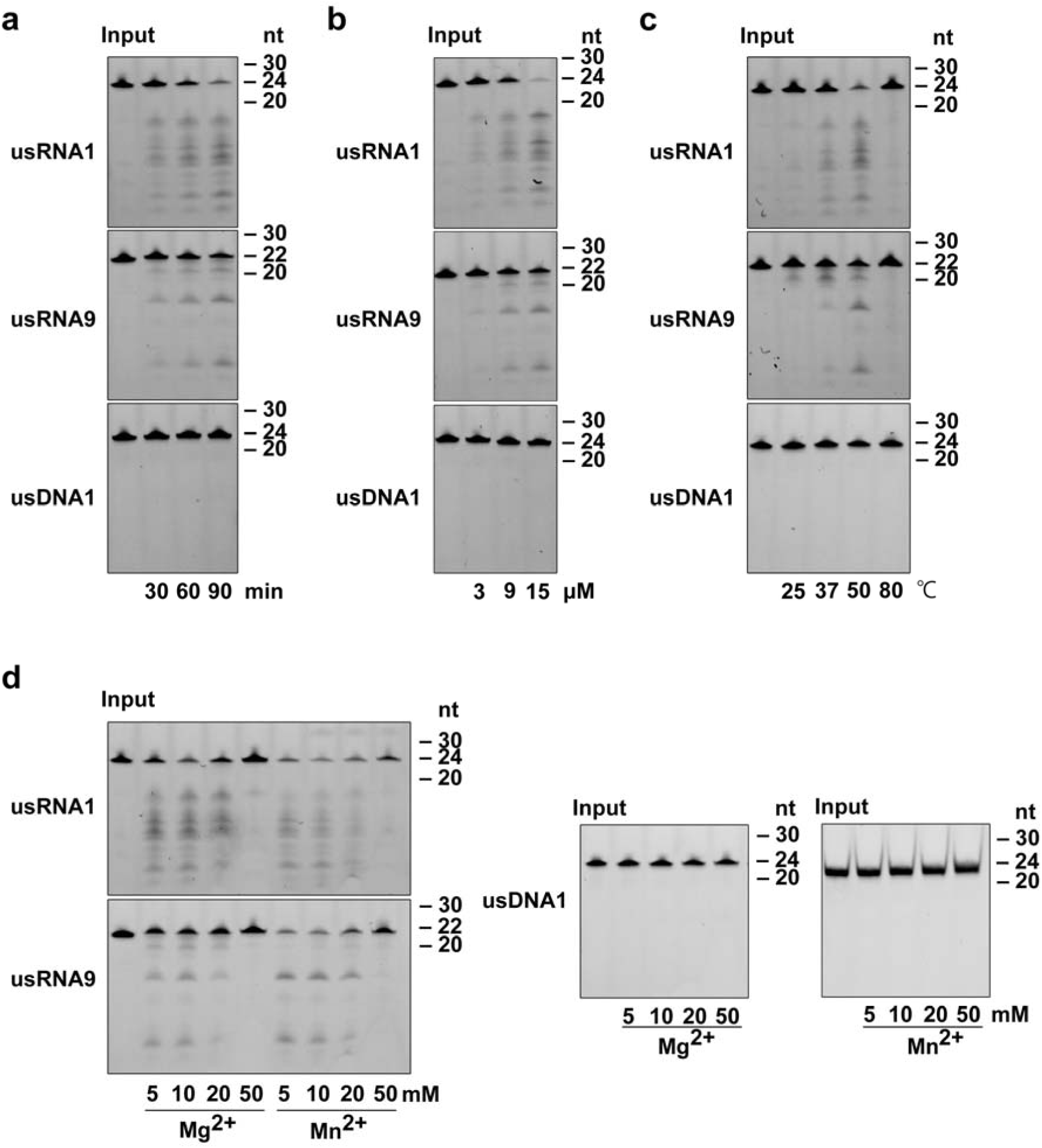
*In vitro* assays for RNase activity of PNGM-1 on small unstructured RNA substrates. The 5′-6-carboxyfluorescein-labeled substrates usRNA1, usRNA9 and usDNA1 were incubated with PNGM-1, and the products were analysed on either a 20% polyacrylamide 8 M urea gel or a 20% polyacrylamide native gel. The native gel was used only for analysis of usDNA1 reactions with MnCl_2_. **a**, The substrates were incubated with 15 μM PNGM-1 in the presence of 10 mM MgCl_2_ at 50 °C for 30, 60 and 90 min. **b**, The substrates were incubated with 3, 9, and 15 μM PNGM-1 in the presence of 10 mM MgCl_2_ at 50 °C for 90 min. **c**, The substrates were incubated with 15 μM PNGM-1 in the presence of 10 mM MgCl_2_ at 25, 37, 50, and 80 °C for 90 min. **d**, The substrates were incubated with 15 μM PNGM-1 in the presence of 5–50 mM MgCl_2_ or MnCl_2_ at 50 °C for 90 min. Experiments in **a-d** were repeated at least three times and were reproducible.

The amounts of usRNA1 and usRNA9 cleavage products increased in a dose-dependent manner, with the optimal temperature for RNase activity estimated to be around 50 °C (Fig. 5b, c). RNase activity was also assessed by varying the concentration (5–50 mM) of MgCl_2_ or MnCl_2_, and the optimal concentrations of Mg^2+^ and Mn^2+^ were estimated to be around 10 mM and 5–10 mM, respectively (Fig. 5d).

### Asp-95 and His-257 are essential for the RNase activity of PNGM-1

To rule out the possibility that the observed RNase activity of PNGM-1 originates from unidentified contaminant RNases, we examined the five PNGM-1 mutants. These mutants contained single amino-acid substitutions of residues that were likely essential for RNase activity on usRNA1 and usRNA9. The PNGM-1 mutants (H91A, H93A, D95A, H96A and H257A: five mutants with alanine replacements of His-48, His-50, Asp-52, His-53 and His-222 at structurally equivalent positions in Tm-tRNase Z, respectively) had a single substitution of alanine for histidine or aspartic acid. The corresponding amino-acid substitutions in Tm-tRNase Z are known to abolish its activity without Mn^2+^ ions. All five PNGM-1 mutants showed little or no RNase activity in the presence of Mg^2+^ (Fig. 6a). These results indicate that the observed RNase activity of PNGM-1 is genuine.

**Fig. 6.**
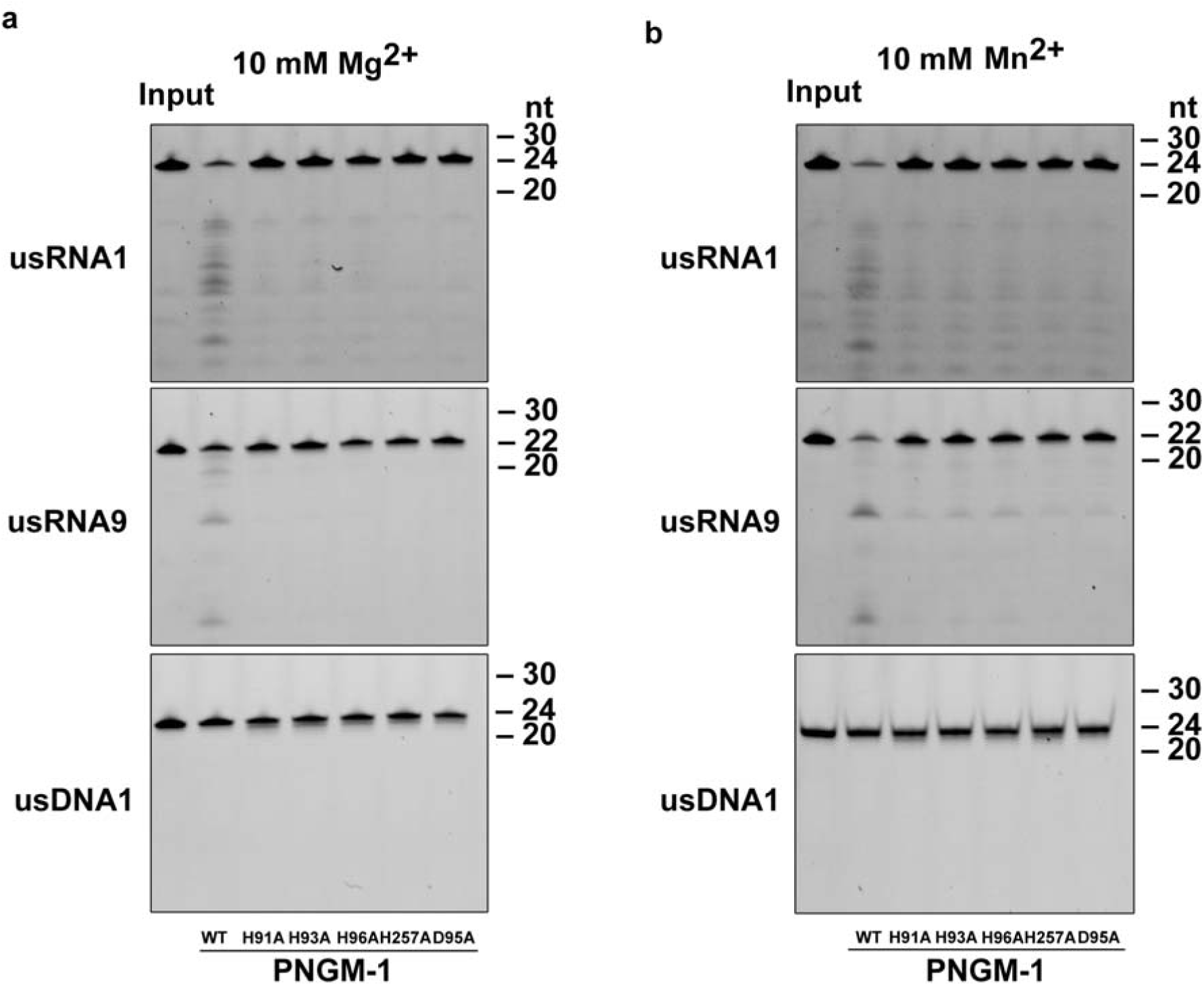
*In vitro* RNA cleavage assays for PNGM-1 mutants. **a**, **b**, usRNA1, usRNA9 and usDNA1 were incubated with 15 μM wild-type PNGM-1 or PNGM-1 mutant (H91A, H93A, H96A, H257A or D95A) in the presence of 10 mM MgCl_2_ (**a**) or MnCl_2_ (**b**), at 50 °C for 90 min, and the products were analysed on a 20% polyacrylamide 8 M urea gel or a 20% polyacrylamide native gel. The native gel was used only for analysis of usDNA1 reactions with MnCl_2_. WT, wild type. Experiments in **a** and **b** were repeated at least three times and were reproducible.

RNase activity was recovered, albeit inefficiently, in the presence of Mn^2+^ for mutants H91A, H93A and H96A but not D95A and H257A (Fig. 6b). This Mn^2+^-rescue phenomenon, which was first observed for the pre-tRNA cleavage reaction by Tm-tRNase Z, suggests that Asp-95 (Asp-52 in Tm-tRNase Z) and His-257 (His-222 in Tm-tRNase Z) are essential and directly contribute to proton transfer for the RNase activity of PNGM-1, further corroborating the RNase activity of PNGM-1. Interestingly, the catalytic efficiency of H257A for all tested β-lactams decreased in comparison to that of wild-type PNGM-1 (Table 1), which suggests that H257 (as well as H91A, H93A, D95A and H96A) of PNGM-1 plays an important role in both its RNase and β-lactamase activities.

### Cleavage site preference of the endoribonucleolytic activity of PNGM-1

We examined the cleavage site preference of the endoribonucleolytic activity of PNGM-1 on usRNA1 and usRNA9. The cleavage reaction products for usRNA1 and usRNA9 were analysed at nt resolutions, with corresponding substrate ladders, on a 20% polyacrylamide denaturing gel. The cleavage patterns of usRNA1 and usRNA9 by PNGM-1 were unique and different from those for Tm-tRNase Z (Fig. 7). PNGM-1 appears to have a tendency to cleave the RNA substrates between pyrimidine nucleotides. The slightly different cleavage patterns observed for Tm-tRNase Z compared with those in the previous study^26^ could be due to the different assay conditions.

**Fig. 7.**
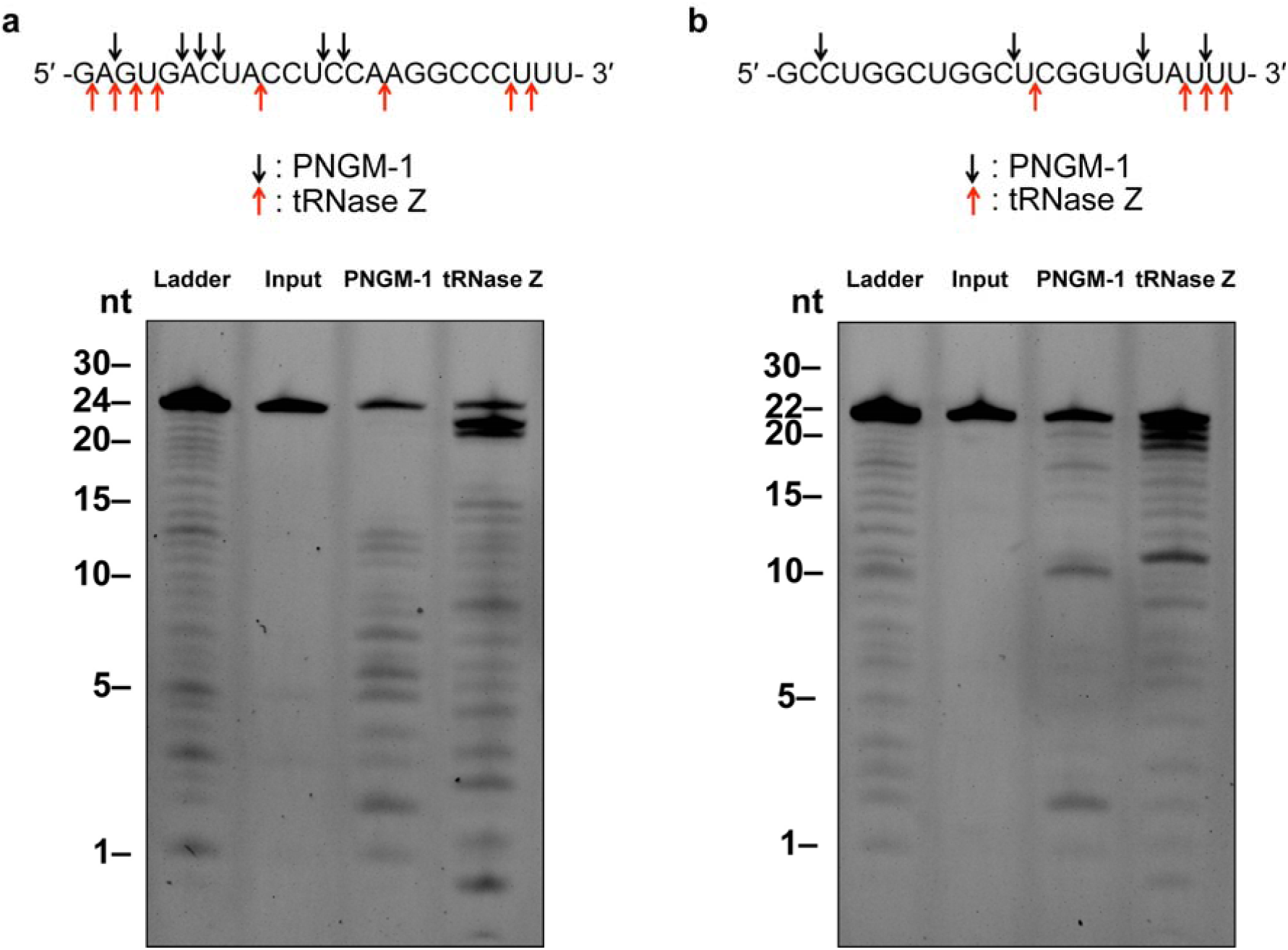
Cleavage site analysis for the RNA substrates usRNA1 and usRNA9. **a**, **b**, usRNA1 (**a**) and usRNA9 (**b**) were incubated at 50 °C in the presence of 10 mM MgCl_2_ with 15 μM PNGM-1 for 90 min or in the presence of 10 mM MnCl_2_ with 1.5 μM Tm-tRNase Z for 30 min. Cleavage products were analysed on a 20% polyacrylamide 8 M urea gel. Arrows indicate the major cleavage sites. Experiments in **a** and **b** were repeated at least three times and were reproducible.

### PNGM-1 can process a pre-tRNA substrate

To test whether PNGM-1 can process pre-tRNA similar to tRNase Z, the 84-nt human pre-tRNA^Arg^, which was 5′-labeled with fluorescein, was used as a pre-tRNA substrate (Fig. 8a). The pre-tRNA^Arg^ was incubated with PNGM-1, Tm-tRNase Z or human Δ30 tRNase Z^L^, with cleavage products analysed by denaturing polyacrylamide gel electrophoresis. PNGM-1 cleaved the pre-tRNA^Arg^ similar to Tm-tRNase Z and human Δ30 tRNase Z^L^ (Fig. 8b). Major cleavage by PNGM-1 occurred after the 76th nt (3 nt downstream of the discriminator), whereas major cleavage by Tm-tRNase Z and human Δ30 tRNase Z^L^ occurred after the 75th nt and the discriminator (73rd nt), respectively, as expected^24,25,27^. Therefore, PNGM-1 is the first unique enzyme that has dual activity i.e., true β-lactamase (MBL hydrolysing all tested and clinically-used β-lactam antibiotics; Table 1) and tRNase Z activity. We are currently trying to co-crystallise PNGM-1 or the PNGM-1 mutants with carbapenems or tRNA^Arg^ in an attempt to understand how PNGM-1 possesses this dual activity.

**Fig. 8.**
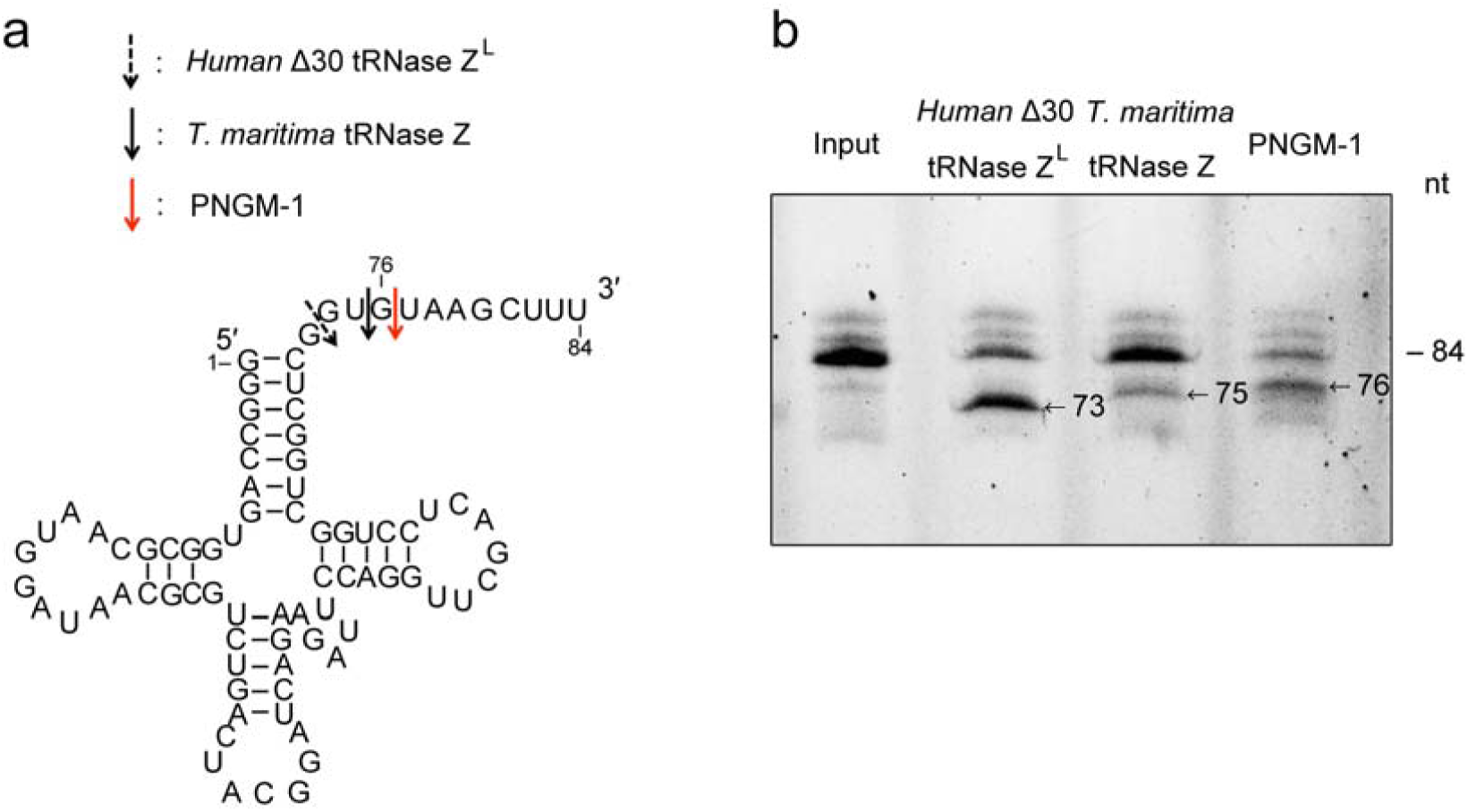
Pre-tRNA processing assay. **a**, The secondary structure of human pre-tRNA^Arg^ and cleavage sites for human Δ30 tRNase Z^L^, Tm-tRNase Z and PNGM-1. **b**, The 84-nt 5′-fluorescein-labeled human pre-tRNA^Arg^ was incubated with 0.5 μM human Δ30 tRNase Z^L^ in the presence of 10 mM MgCl_2_ at 37 °C for 30 min, with 1.5 μM Tm-tRNase Z at 37 °C for 60 min or with 15 μM PNGM-1 at 50 °C for 90 min. The products were analysed on a 20% polyacrylamide 8 M urea gel. Experiments in **a** and **b** were repeated at least three times and were reproducible.

### Origin of subclass B3 MBLs

Our functional analyses of PNGM-1 showed that PNGM-1 has true β-lactamase (MBL) and tRNase Z activity (Table 1 and Figs. 3-7). We demonstrate the dual activity of PNGM-1, which strongly suggests that PNGM-1 has evolved from a tRNase Z [MBL fold protein with a single native activity and without true-β-lactamase activity (Supplementary Table S5)]. This is in agreement with earlier reports suggesting that novel enzymes could evolve from an existing enzyme with single native activity through the recruitment of multiple functions (e.g., dual activity)^20,36^. It is also consistent with previous reports showing that (i) MBLs descended from a common ancestor (MBL fold protein) of the MBL superfamily^17^ and (ii) the last common ancestor of the MBL superfamily is a phosphodiesterase, such as Ec-tRNase Z (PDB code: 2CBN), involved in tRNA processing^20,37^. In addition, comparison of the PNGM-1 structure with all structures in the Protein Data Bank, using DALI server (http://ekhidna2.biocenter.helsinki.fi/dali/), revealed that the highest structural similarity is for Bs-tRNase Z (Dali Z-score: 29.7), followed by Ec-tRNase Z (29.4), Tm-tRNase Z (22.2) and subclass B3 MBLs [AIM-1 (10.0), GOB-18 (9.8) and FEZ-1 (9.7)] (Supplementary Table S6). Analysis of the superimposed structures of these six enzymes with PNGM-1 showed the gradual change from tRNase Z to subclass B3 MBL (Fig. 2). tRNA is a bulky substrate similar to tRNase Z enzyme monomer. In order to recognize tRNA, tRNase Z has an elongated dimeric structure with an extended tRNA-binding arm protruding from the central β-sheet in the αββα-fold (Fig. 2). In the dimer structure, subunit A is responsible for substrate recognition while subunit B is catalytically active (Fig. 2). The tRNA-binding arm has a positively charged surface which provides an extra binding affinity for the negatively charged tRNA phosphate backbone (Supplementary Fig. S1). PNGM-1 has maintained the dimerizing ability but has lost the tRNA-binding arm. Subclass B3 MBLs has lost the dimerization. However, the dimerization of subclass B3 MBLs is not necessary for the binding of much smaller β-lactam substrates. When tRNA was modelled to bind to subclass B3 MBLs, subclass B3 MBLs are unable to interact with tRNA except the 3’ end of tRNA to be processed (Fig. 2). Therefore, among subclass B3 MBLs, only PNGM-1 has true tRNase Z activity.

We compared the PNGM-1 structure with carbapenem-bound subclass B3 MBL structures such as diapenem-bound CphA and doripenem-bound SMB-1 (Supplementary Fig. S2). The superimposed diapenem and doripenem are almost in the same position as the 3’ end of tRNA close to the metal ions in the active site (Figs. 2 and 3a). The substrate-binding pocket between PNGM-1 and subclass B3 MBLs is well conserved (Figs. 3b-d). Furthermore, the efficiency (*k*_cat_/*K*_*m*_ for carbapenems [imipenem, meropenem and ertapenem]) of PNGM-1 MBL activity is significantly lower than that of the single native MBL activity of AIM-1, GOB-18 or FEZ-1 by 3 or 4 orders (Table 1, Supplementary Table S5) and lower than that of Tm-tRNase Z by 3 orders^24^. β-Lactam-containing antibiotics including carbapenems (substrates for MBLs) existed in natural environments^8^ and natural antibiotics appear to exist in the biosphere billions years ago^38-40^. It means that the B3 MBL activity of PNGM-1 is a promiscuous activity and subclass B3 MBLs is thought to have evolved from PNGM-1. It is consistent with the Jensen conceptualised enzyme evolution showing that newly specialised enzymes are generated from promiscuous activities^20,41^.

Based on our results, we suggest that subclass B3 MBLs arose through an evolutionary process (tRNase Z → PNGM-1→ subclass B3 MBLs) and the origin of subclass B3 MBLs is a tRNase Z. If a B1/B2 MBL with dual activity, like PNGM-1, could be identified in the future, we would be able to understand the evolutionary origin of subclass B1/B2 MBLs, as MBLs have evolved independently twice (B1/B2 and B3 subclasses) billions years ago^17,20,42^. These evolutionary processes help us to understand where MBL genes came from and predict the future evolution of MBL genes, as previously described^40^.

## Acknowledgements

We thank our colleagues, both past and present, at the National Leading Research Laboratory of Drug Resistance Proteomics, Myongji University, for sharing their insights into the concepts presented here; S.G. Kang and B.C. Jeong for early support of this work; and J.H. Kim and S.K. Malik for helpful discussions. This work was supported by research grants from the Bio & Medical Technology Development Program of the National Research Foundation of Korea (NRF) funded by the Ministry of Science and ICT (MSIT; grants NRF-2017M3A9E4078014 and NRF-2017R1A2B4002315 to S.H.L. NRF-2017M3A9E4078017 to L.-W.K., and NRF-2019R1C1C1008615 to J.H.L.); the Korea Centers for Disease Control and Prevention (grant 2017ER540402 to S.H.L.); and the Takeda Science Foundation to M.T.

## Author contributions

J.H.L., M.T., J.H.J. and L.-W.K. contributed equally. J.H.L., J.H.J and J.-H.L. identified and prepared a novel MBL gene and protein. J.H.L., M.T., M.N., K.S.P. and M.S. performed a genetic and functional analysis. J.H.J., T.Y.K. and J.H.K. conducted the mutant analysis. L.-W.K., M.-K.H. and Y.S.P. carried out the crystal structure studies and data analysis. S.H.L. and M.N. analysed the evolutionary origin. All authors contributed to manuscript preparation. S.H.L. provided overall guidance for the design and execution of this work.

## Additional information

### Correspondence and requests for materials

should be addressed to M.N. or S.H.L.

