## Supplemental Tables S1-S6 and Figures S1-S2 for "Dual activity of PNGM-1, a metallo-β-lactamase and tRNase Z, pinpoints the evolutionary origin of subclass B3 metallo-β-lactamases"

**Subtitle: Origin of subclass B3 metallo-β-lactamases**

Jung Hun Lee[1,5], Masayuki Takahashi[2,5], Jeong Ho Jeon[1,5], Lin-Woo Kang[3,5], Mineaki Seki[2], Kwang Seung Park[1], Myoung-Ki Hong[3], Yoon Sik Park[3], Tae Yeong Kim[1], Asad Mustafa Karim[1], Jung-Hyun Lee[4], Masayuki Nashimoto[2,6]* & Sang Hee Lee[1,6]*

[1]National Leading Research Laboratory of Drug Resistance Proteomics, Department of Biological Sciences, Myongji University, 116 Myongjiro, Yongin, Gyeonggido 17058, Republic of Korea. [2]Research Institute for Healthy Living, Niigata University of Pharmacy and Applied Life Sciences, Higashijima 265-1, Niigata, Niigata 956-8603, Japan. [3]Department of Biological Sciences, Konkuk University, 1 Hwayang-dong, Gwangjin-gu, Seoul 05029, Republic of Korea. [4]Marine Biotechnology Research Center, Korea Institute of Ocean Science & Technology, 385 Haeyang-ro, Yeongdo-gu, Busan 49111, Republic of Korea.

[5]These authors contributed equally: Jung Hun Lee, Masayuki Takahashi, Jeong Ho Jeon, Lin-Woo Kang.
[6]These authors jointly supervised this work: Masayuki Nashimoto, Sang Hee Lee.
*

**Supplementary Table S1 | Strains and plasmids used in this study**

| Strains and plasmids | Phenotype, genotype and/or characteristics | Source (or reference) |
|---|---|---|
| Strains | | |
| *E. coli* BL21(DE3) | $F^-$ *ompT hsdS*$_B$($r_B^- m_B^-$) *gal dcm* (DE3) | Invitrogen |
| Plasmids | | |
| pET-28a(+) | Expression vector, kanamycin$^r$ | Novagen |
| pET-30a(+) | Expression vector, kanamycin$^r$ | Novagen |
| pET-28a(+)/His$_6$-PNGM-1 | The *bla*$_{PNGM-1}$ gene was cloned into the pET-28a(+) vector, encoding N-terminal His$_6$-tag | Park *et al.* (2018)[12] |
| pET-28a(+)/His$_6$-PNGM-1 (H91A) | H91 in PNGM-1 was replaced by alanine by site-directed mutagenesis | This study |
| pET-28a(+)/His$_6$-PNGM-1 (H93A) | H93 in PNGM-1 was replaced by alanine by site-directed mutagenesis | This study |
| pET-28a(+)/His$_6$-PNGM-1 (D95A) | D95 in PNGM-1 was replaced by alanine by site-directed mutagenesis | This study |
| pET-28a(+)/His$_6$-PNGM-1 (H96A) | H96 in PNGM-1 was replaced by alanine by site-directed mutagenesis | This study |
| pET-28a(+)/His$_6$-PNGM-1 (H257A) | H257 in PNGM-1 was replaced by alanine by site-directed mutagenesis | This study |
| pET-30a(+)/His$_6$-AIM-1 | The *bla*$_{AIM-1}$ gene was cloned into the pET-30a(+) vector | This study |
| pET-30a(+)/His$_6$-GOB-18 | The *bla*$_{GOB-18}$ gene was cloned into the pET-30a(+) vector | This study |
| pET-30a(+)/His$_6$-FEZ-1 | The *bla*$_{FEZ-1}$ gene was cloned into the pET-30a(+) vector | This study |
| pET-30a(+)/His$_6$-Bs-tRNase Z | The tRNase Z gene from *Bacillus subtilis* was cloned into the pET-30a(+) vector | This study |
| pET-30a(+)/His$_6$-Ec-tRNase Z | The tRNase Z gene from *Escherichia coli* was cloned into the pET-30a(+) vector | This study |
| pET-30a(+)/His$_6$-Tm-tRNase Z | The tRNase Z gene from *Thermotoga maritima* was cloned into the pET-30a(+) vector | This study |

r: resistant.

**Supplementary Table S2 | Primers used in this study**

| Name | Sequence (5′→3′) |
|---|---|
| Primers for site-directed mutagenesis | |
| PNGM-1-H91A-F[a] | 5'–AAGATTTTTCTGACGGCCTTGCACACCGACCAC–3' |
| PNGM-1-H91A-R[b] | 5'–GTGGTCGGTGTGCAAGGCCGTCAGAAAAATCTT–3' |
| PNGM-1-H93A-F | 5'–TTTCTGACGCATTTGGCCACCGACCACTGGGGC–3' |
| PNGM-1-H93A-R | 5'–GCCCCAGTGGTCGGTGGCCAAATGCGTCAGAAA–3' |
| PNGM-1-D95A-F | 5'–ACGCATTTGCACACCGCCCACTGGGGCGACCTG–3' |
| PNGM-1-D95A-R | 5'–CAGGTCGCCCCAGTGGGCGGTGTGCAAATGCGT–3' |
| PNGM-1-H96A-F | 5'–CATTTGCACACCGACGCCTGGGGCGACCTGGTG–3' |
| PNGM-1-H96A-R | 5'–CACCAGGTCGCCCCAGGCGTCGGTGTGCAAATG–3' |
| PNGM-1-H257A-F | 5'–ATCAATCTGGACTTTGCCACCTCAGCGCAATCC–3' |
| PNGM-1-H257A-R | 5'–GGATTGCGCTGAGGTGGCAAAGTCCAGATTGAT–3' |
| Primers for cloning | |
| *Nde*I-His-EK-AIM-1-F | 5'–ATA**CATATGCATCATCATCATCATCAT***GACGACGACGACAAG*<br>TCGGATGCTCCAGCCTCAAGAGGA–3' |
| *Xho*I-AIM-1-R | 5'–GAG**CTCGAG**TCAAGGCCGAGCACCACTAGAC–3' |
| *Nde*I-His-EK-GOB-18-F | 5'–ATA**CATATGCATCATCATCATCATCAT***GACGACGACGACAAG*<br>GCTCAGGTAGTAAAAGAACCTGAAAATAT–3' |
| *Xho*I- GOB-18-R | 5'–CAG**CTCGAG**TTATTTCTTTATTGCATTCAGCACAG–3' |
| *Nde*I-His-EK-FEZ-1-F | 5'–ATA**CATATGCATCATCATCATCATCAT***GACGACGACGACAAG*<br>GCTTATCCAATGCCTAACCCC–3' |
| *Xho*I-FEZ-1-R | 5'–CAG**CTCGAG**TTATTTATCTTGGGAATCTTTTTTTATTTTGTTGA<br>GAT–3' |
| *Nde*I-His-EK-Bs-tRNaseZ-F | 5'–ATA**CATATGCATCATCATCATCATCAT***GACGACGACGACAAG*<br>GAGTTATTATTCTTGGGTACTGGTGCGG–3' |
| *Xho*I-Bs-tRNaseZ-R | 5'–GAG**CTCGAG**TCAACCGCGGGGAACGTTGACT–3' |
| *Nde*I-His-EK-Ec-tRNaseZ-F | 5'–ATA**CATATGCATCATCATCATCATCAT***GACGACGACGACAAG*<br>GAATTAATTTTTTTAGGTACTTCAGCCGG–3' |
| *Xho*I-Ec-tRNaseZ-R | 5'–CAG**CTCGAG**TTAAACGTTAAACACGGTGAAATCATTCGCC–3' |
| *Nde*I-His-EK-Tm-tRNaseZ-F | 5'–ATA**CATATGCATCATCATCATCATCAT***GACGACGACGACAAG*<br>AACATAATCGGCTTCAGCAAAG –3' |
| *Xho*I-Tm-tRNaseZ-R | 5'–GAG**CTCGAG**TCACATTTCAAATACTTTTCTCGGG–3' |

The positions of the mutated codons are underlined. Restriction sites appear in bold. The underlined and bolded bases indicate the hexahistidine tag, and italic bases indicate the enterokinase recognition site.

[a]F, forward.

[b]R, reverse.

**Supplementary Table S3 | List of the representative types of metallo-β-lactamases (MBLs) and structurally representative MBL fold proteins used to construct the phylogenetic trees for PNGM-1**

| Subclass B1 MBLs | | Subclasses B2 and B3 MBLs | | MBL fold proteins | | |
|---|---|---|---|---|---|---|
| Name | Accession no.[a] | Name | Accession no.[a] | Name | Accession no.[a] or UniProt ID[b] | Description |
| ANA-1 | WP_041449074 | CphA-1 | CAA40386 | Bs-tRNase Z | P54548[b] | tRNase Z or Ribonuclease Z |
| BclI-1 | AAA22276 | SFH-1 | WP_024531368 | Tm-tRNase Z | AKE26778[a] | |
| BlaB-1 | AAF89154 | ImiS | CAA71441 | Ec-tRNase Z | P0A8V0[b] | |
| CfiA | AAA22907 | ImiH | CAD69003 | Tm | NP_228022[a] | Zn-dependent hydrolase |
| CGB-1 | AAL55263 | PNGM-1 | AWN09461 | Tm-1 | Q9X207[b] | Zn-dependent hydrolase |
| DIM-1 | AGC92784 | AIM-1 | CAQ53840 | Tm-Lac | Q9WZZ6[b] | Lactonase |
| EBR-1 | AAN32638 | ALG6-1 | APR64488 | AiiA | P0CJ63[b] | N-Acyl Homoserine Lactone Hydrolase |
| ECV-1 | AGA78874 | ALG11-1 | APR64489 | PDLA | Q988B9[b] | 4-pyridoxolactonase |
| ElBla2-1 | ABC63608 | BJP-1 | BAC51495 | SdsA1 | AAG04129[a] | Alkylsulfatase |
| FIA-1 | WP_041258349 | CAR-1 | AIA71664 | ATSD | Q9C8L4[b] | Sulfur dioxygenase |
| FIM-1 | AFV91534 | CAU-1 | CAC87665 | CbpE | CAC29434[a] | Teichoic acid phosphorylcholine esterase |
| GIM-1 | ALO69078 | CPS-1 | AJP77054 | Pah | AAP06948[a] | Methyl parathion hydrolase |
| GRD23-1 | APR64493 | CRD3-1 | APR64487 | Gox | Q16775[b] | Glyoxalase II |
| HMB-1 | AMY61250 | DHT2-1 | APR64485 | MTH1203 | WP_010876827[a] | β-CASP metallo-β-lactamase family nuclease |
| IMP-1 | ABK27309 | EAM-1 | AFN85388 | | | |
| IND-1 | AAD20273 | ECM-1 | AFN85387 | | | |
| JOHN-1 | AAK38324 | EFM-1 | AFN85384 | | | |
| KHM-1 | BAF91108 | ELM-1 | AFN85386 | | | |
| MOC-1 | ANJ59787 | ESP-1 | AJP77085 | | | |
| MUS-1 | AAN63647 | EVM-1 | AFN85385 | | | |
| MYO-1 | WP_081048762 | FEZ-1 | CAB96921 | | | |
| MYX-1 | ABF86854 | GOB-1 | AAF04458 | | | |
| NDM-1 | AHM26723 | L1-1 | CAA52968 | | | |
| ORR-1 | WP_109545042 | LRA3-1 | ACH58987 | | | |
| PEDO-3 | AJP77076 | LRA7-1 | ACH58998 | | | |
| PST-1 | WP_043942497 | LRA8-1 | ACH58988 | | | |
| SFB-1 | AAT90847 | LRA12-1 | ACH58990 | | | |
| SHD-1 | ABE54111 | LRA17-1 | ACH58994 | | | |
| SHN-1 | ABE56430 | LRA19-1 | ACH59005 | | | |
| SIM-1 | AER61546 | LRA2-1 | ACH58985 | | | |
| SLB-1 | AAT90846 | MEMA1-1 | KY705336 | | | |
| SPM-1 | AAR15341 | MIM-1 | AIT78529 | | | |
| SPN79-1 | APR64486 | MSI-1 | AJP77057 | | | |
| SPS-1 | ADK81930 | PEDO-1 | AJP77059 | | | |
| STA-1 | WP_109545039 | PLN-1 | KIO75746 | | | |
| TTU-1 | ACR12883 | POM-1 | ABY56045 | | | |
| TMB-1 | CBY88906 | RM3 | AGU01679 | | | |
| TUS-1 | EKB08120 | SAG-1 | AFV00127 | | | |
| VIM-1 | CAC35170 | SMB-1 | BAL14456 | | | |
| ZOG-1 | CAZ94871 | SPG-1 | AJP77080 | | | |
| | | SPR-1 | ABV42357 | | | |
| | | THIN-B | CAC33832 | | | |

[a]NCBI GenBank database accession number (http://www.ncbi.nlm.nih.gov/).
[b]UniProt is a freely accessible database of protein sequence and functional information (https://www.uniprot.org/).

**Supplementary Table S4 | Data collection and refinement statistics**

| **Data collection** | Native | SeMet |
|---|---|---|
| Space group | $P2_1$ | $P2_1$ |
| Unit-cell parameters | | |
| a, b, c (Å) | 122.3, 83.0, 163.5 | 121.9, 83.1, 162.8 |
| α, β, γ (°) | 90, 110.6, 90 | 90, 110.2, 90 |
| Resolution (Å) | 50.0 – 2.1 (2.14 – 2.10) | 50.0 – 2.3 (2.34 – 2.30) |
| Total reflections | 1,318,149 | 1,018,484 |
| Unique reflections | 175,405 | 135,186 |
| Completeness (%) | 98.2 (97.5) | 98.1 (86.8) |
| Multiplicity | 7.5 (7.4) | 7.5 (6.9) |
| $\langle I/\sigma(I) \rangle$ | 47.7 (15.5) | 46.8 (11.3) |
| $R_{merge}$ (%) | 9.0 (22.7) | 12.2 (34.2) |
| **Refinement** | | |
| Resolution (Å) | 34.3–2.1 (2.17–2.10) | 33.3-2.3 (2.37-2.29) |
| Used reflections | 175,021 (17,252) | 134,467 (12,544) |
| Macromolecules / asymmetric unit | 8 | 8 |
| $R_{work}$ / $R_{free}$ (%) | 19.2/24.3 | 21.3/26.8 |
| No. of atoms | 24,786 | 23,910 |
| Protein | 23,464 | 23,464 |
| Water | 1,306 | 430 |
| $Zn^{2+}$ ion | 16 | 16 |
| B-factor | 25.5 | 32.9 |
| Protein | 25.5 | 33.0 |
| Water | 25.2 | 26.2 |
| $Zn^{2+}$ ion | 24.1 | 31.4 |
| RMSD | | |
| Bond length (Å) | 0.019 | 0.016 |
| Bond angles (º) | 1.85 | 1.72 |
| Ramachandran plot (%) | | |
| Favored | 95.0 | 94.0 |
| Allowed | 4.1 | 5.0 |
| Disallowed | 0.9 | 1.0 |

Dataset was collected from a single crystal. Data collection data was previously published[16]. Values in parentheses are for the shell with the highest-resolution. $R_{merge} = \sum_{hkl}\sum_i \left|\left(I_i(hkl)\right) - \langle I(hkl)\rangle\right| / \sum_{hkl}\sum_i I_i(hkl)$, where $I_i(hkl)$ is the mean intensity of the $i$th observation of symmetry-related reflections *hkl*. $R_{work} = \sum_{hkl}\left||F_{obs}| - |F_{calc}|\right| / \sum_{hkl}|F_{obs}|$, where $F_{calc}$ is the calculated protein structure factor from the atomic model ($R_{free}$ was calculated as $R_{work}$ with a randomly selected 5% of the reflections). Experiments were repeated at least three times and were reproducible.

**Supplementary Table S5 | Kinetic parameters of AIM-1, GOB-18, FEZ-1 and three tRNase Zs for various β-lactams**

| Substrate and parameter | AIM-1 | GOB-18 | FEZ-1 | Ec-tRNase Z | Bs-tRNase Z | Tm-tRNase Z |
|---|---|---|---|---|---|---|
| Cefoxitin | | | | | | |
| $K_m$ (μM) | 10.1 ± 0.1 | 12.5 ± 0.1 | 9.2 ± 0.1 | NH | NH | NH |
| $k_{cat}$ ($s^{-1}$) | 4.725 ± 0.001 | 2.628 ± 0.001 | 1.564 ± 0.001 | NH | NH | NH |
| $k_{cat}/K_m$ ($M^{-1}$ $s^{-1}$) | (4.7 ± 0.2) × $10^5$ | (2.1 ± 0.2) × $10^5$ | (1.7 ± 0.1) × $10^5$ | NH | NH | NH |
| Ceftazidime | | | | | | |
| $K_m$ (μM) | 8.3 ± 0.2 | 10.3 ± 0.2 | 7.6 ± 0.2 | NH | NH | NH |
| $k_{cat}$ ($s^{-1}$) | 3.718 ± 0.001 | 2.067 ± 0.001 | 3.511 ± 0.001 | NH | NH | NH |
| $k_{cat}/K_m$ ($M^{-1}$ $s^{-1}$) | (4.5 ± 0.2) × $10^5$ | (2.0 ± 0.2) × $10^5$ | (4.6 ± 0.2) × $10^5$ | NH | NH | NH |
| Cefotaxime | | | | | | |
| $K_m$ (μM) | 5.4 ± 0.1 | 6.7 ± 0.1 | 4.9 ± 0.1 | NH | NH | NH |
| $k_{cat}$ ($s^{-1}$) | 2.157 ± 0.002 | 1.2 ± 0.003 | 1.205 ± 0.002 | NH | NH | NH |
| $k_{cat}/K_m$ ($M^{-1}$ $s^{-1}$) | (4.0 ± 0.1) × $10^5$ | (1.8 ± 0.1) × $10^5$ | (2.5 ± 0.1) × $10^5$ | NH | NH | NH |
| Meropenem | | | | | | |
| $K_m$ (μM) | 4.6 ± 0.1 | 5.7 ± 0.1 | 4.2 ± 0.1 | NH | NH | NH |
| $k_{cat}$ ($s^{-1}$) | 15.409 ± 0.0001 | 8.568 ± 0.001 | 0.773 ± 0.001 | NH | NH | NH |
| $k_{cat}/K_m$ ($M^{-1}$ $s^{-1}$) | (3.3 ± 0.1) × $10^6$ | (1.5 ± 0.1) × $10^6$ | (1.8 ± 0.1) × $10^5$ | NH | NH | NH |
| Imipenem | | | | | | |
| $K_m$ (μM) | 3.8 ± 0.1 | 4.8 ± 0.1 | 3.5 ± 0.1 | NH | NH | NH |
| $k_{cat}$ ($s^{-1}$) | 11.128 ± 0.0001 | 6.188 ± 0.003 | 0.763 ± 0.001 | NH | NH | NH |
| $k_{cat}/K_m$ ($M^{-1}$ $s^{-1}$) | (2.9 ± 0.2) × $10^6$ | (1.3 ± 0.2) × $10^6$ | (2.2 ± 0.1) × $10^5$ | NH | NH | NH |
| Ertapenem | | | | | | |
| $K_m$ (μM) | 5.0 ± 0.1 | 6.3 ± 0.1 | 4.6 ± 0.1 | NH | NH | NH |
| $k_{cat}$ ($s^{-1}$) | 12.376 ± 0.0001 | 6.882 ± 0.002 | 0.667 ± 0.001 | NH | NH | NH |
| $k_{cat}/K_m$ ($M^{-1}$ $s^{-1}$) | (2.5 ± 0.2) × $10^6$ | (1.1 ± 0.2) × $10^6$ | (1.5 ± 0.2) × $10^5$ | NH | NH | NH |

[a]NH, not hydrolysed.

Data are mean ± s.d. of three assays.

**Supplementary Table S6 | Comparison of PNGM-1 structure against all structures in the Protein Data Bank (PDB)**

| Organism | Chain | Z[a] | rmsd[b] | lali[c] | nres[d] | %id[e] | Protein |
|---|---|---|---|---|---|---|---|
| *Bacillus subtilis* | 4GCW-A | 29.7 | 2.2 | 251 | 307 | 20 | Bs-tRNase Z |
| *Escherichia coli* | 2CBN-A | 29.4 | 2 | 249 | 306 | 21 | Ec-tRNase Z |
| *Thermotoga maritima* | 2E7Y-A | 22.2 | 2.5 | 220 | 272 | 16 | Tm-tRNase Z |
| *Pseudomonas aeruginosa* | 4AWZ-A | 10.0 | 3.7 | 166 | 269 | 13 | AIM-1 |
| *Elizabethkingia meningosep* | 5K0W-A | 9.8 | 3.3 | 163 | 270 | 13 | GOB-18 |
| *Legionella gormanii* | 5W90-A | 9.7 | 3.2 | 162 | 263 | 8 | FEZ-1 |

A total of 883 hits were found from the DALI search. Bs-tRNase Z had the highest Z-score (29.7) and ArcA had the lowest Z-score (2.0). Similarities with Z-scores lower than 2 are spurious.
[a]Z score, the statistical significance of the similarity between the protein-of-interest and other neighboring proteins.
[b]Root Mean Square Distance (rmsd), root-mean-square deviation of C-alpha atoms in the least-squares superimposition of the structurally equivalent C-alpha atoms.
[c]lali, the number of structurally equivalent residues.
[d]nres, the total number of amino acids in the hit protein.
[e]%id, the percentage of identical amino acids over structurally equivalent residues.
Representative comparison results are shown in this Table.

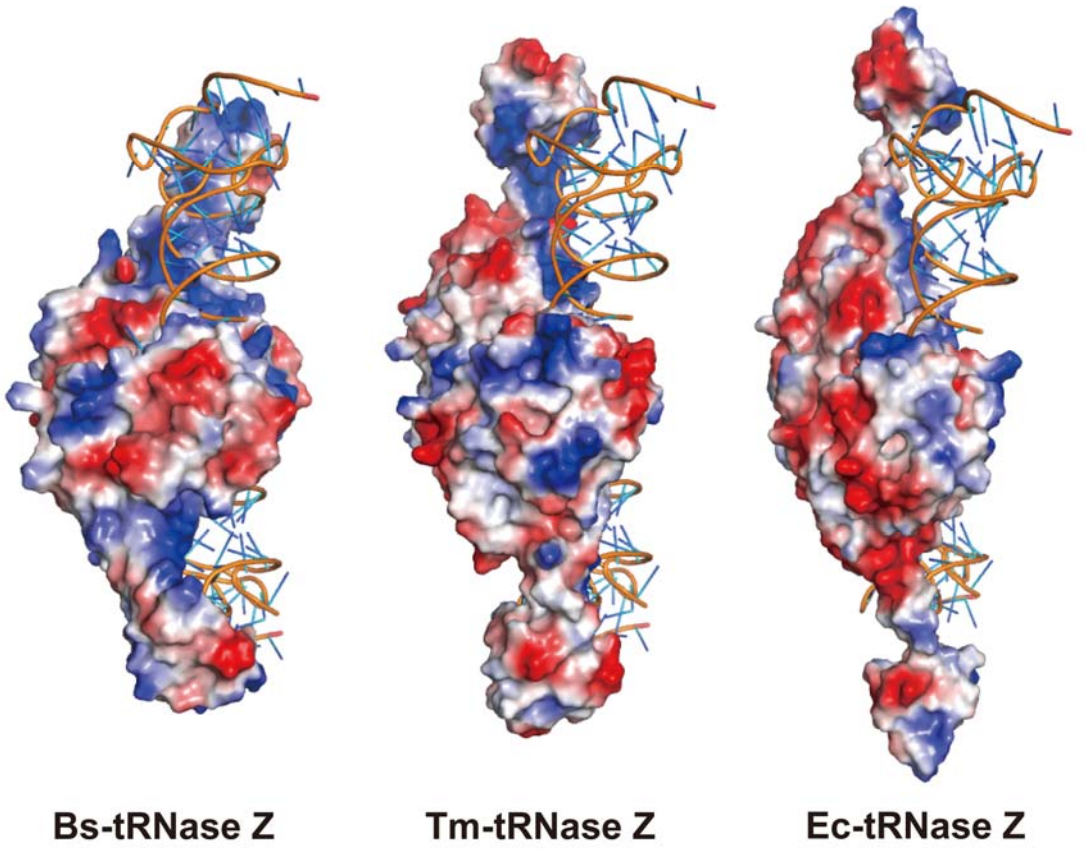

**Supplementary Fig. S1 |** The surface electrostatic potential of Bs-tRNase Z (co-crystallised with tRNA, PDB entry 4GCW) (**a**), Tm-tRNase Z (with superimposed tRNA, PDB entry 2E7Y) (**b**), and Ec-tRNase Z (with superimposed tRNA, PDB entry 2CBN) (**c**). Positively and negatively charged regions are shown in blue and red, respectively.

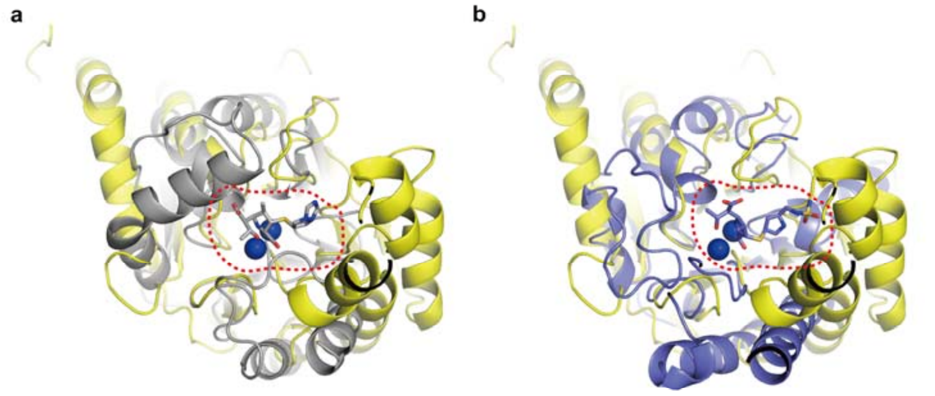

**Supplementary Fig. S2 |** Structure comparison of PNGM-1 with a carbapenem-bound MBL. **a, b,** PNGM-1 (yellow) is superimposed with diapenem-bound CphA (grey, PDB entry 1X8I) (**a**) and doripenem-bound SMB-1 (pale purple, PDB entry 5B15) (**b**). The two metal ions in the active site are shown (blue sphere). The substrate-binding pocket is marked with a red dashed line.
